# Neurodegeneration Risk Factor, *EIF2AK3* (*PERK*), Influences Tau Protein Aggregation

**DOI:** 10.1101/2022.12.14.520487

**Authors:** Goonho Park, Ke Xu, Leon Chea, Kyle Kim, Lance Safarta, Keon-Hyoung Song, Jian Wu, Soyoung Park, Hyejung Min, Nobuhiko Hiramatsu, Jaeseok Han, Jonathan H. Lin

## Abstract

Tauopathies are neurodegenerative diseases caused by pathologic misfolded tau protein aggregation in the nervous system. Population studies implicate *EIF2AK3* (*eukaryotic translation initiation factor 2 alpha kinase 3*), better known as *PERK* (*protein kinase R-like endoplasmic reticulum kinase*), as a genetic risk factor in several tauopathies. PERK is a key regulator of intracellular proteostatic mechanisms – Unfolded Protein Response (UPR) and Integrated Stress Response (ISR). Previous studies found that tauopathy-associated PERK variants encoded functional hypomorphs with reduced signaling *in vitro*. But, it remained unclear how altered PERK activity led to tauopathy. Here, we chemically or genetically modulated PERK signaling in cell culture models of tau aggregation and found that PERK pathway activation prevented tau aggregation while inhibition exacerbated tau aggregation. In primary tauopathy patient brain tissues, we found that reduced PERK signaling correlated with increased tau neuropathology. We found that tauopathy-associated PERK variants targeted the ER luminal domain; and two of these variants damaged hydrogen bond formation. Our studies support that PERK activity protects against tau aggregation and pathology. This may explain why people carrying hypomorphic PERK variants have increased risk for developing tauopathies. Finally, our studies identify small molecule augmentation of PERK signaling as an attractive therapeutic strategy to treat tauopathies by preventing tau pathology.

## Introduction

Tauopathies are age-related neurodegenerative diseases that include Alzheimer’s disease (AD) and Progressive Supranuclear Palsy (PSP) (1–4). Different brain regions are affected in these diseases that account for varying clinical presentations, but tauopathies all lead to progressive, irreparable morbidity that can quickly progress to mortality. In people, the microtubule-associated protein tau (*MAPT)* gene encodes tau protein and is abundantly transcribed throughout the brain (5–8). Alternative splicing of the *MAPT* transcript generates 6 tau protein isoforms that carry varying numbers of carboxy-terminal repeat (R) domains (5,9,10). In healthy cells, tau stabilizes and regulates microtubule assembly and is highly enriched in axons, and also found in dendrites, nuclei, and extracellular space (5,8,11). By contrast, in tauopathies, tau adopts abnormal conformations, becomes hyperphosphorylated, and forms dense aggregates in neurons (2,5,7,12). Environmental and genetic risk factors have been identified that influence tauopathy disease development and progression, but their pathomechanisms are incompletely understood (4).

*EIF2AK3*, more commonly known as PERK, is a genetic risk factor for tauopathies: PSP(13–15) and AD(16,17). PERK is an important regulator of the UPR and ISR(18–20). In response to ER stress protein misfolding, PERK slows cellular translation by phosphorylating eukaryotic initiation factor 2 alpha (eIF2*α*)(18). PERK signaling also initiates a characteristic transcriptional program through induction of transcription factors including ATF4, which upregulates GADD34 phosphatase, and CHOP (21–23). The GADD34 converts P-eIF2*α*to eIF2*α* and thereby restores translation (24). *PERK^-/-^* mice develop marked endocrine and exocrine pancreatic cell death leading to diabetes mellitus(25), and this phenotype is closely recapitulated in Wolcott-Rallison syndrome (WRS), an autosomal recessive genetic disease caused by variants in human PERK(26,27). Tauopathy symptoms are not features of WRS. Conversely, diabetes and/or pancreatic insufficiency are not primary features of tauopathies. It is unclear why PERK is linked with such markedly different human diseases.

Previously, we found that tauopathy-risk *PERK* variants showed reduced protein stability and signaling compared to protective *PERK* variants in cell culture assays(28). We also found that iPSC-derived neurons from tauopathy patients showed reduced phosphorylation of eIF2*α* when challenged with ER stress-inducing chemicals(28). Based on these findings, we proposed that tauopathy-associated PERK variants are functional hypomorphs and that changes in PERK signaling somehow influence the development of these neurodegenerative diseases. Here, we further evaluated the role of PERK in tauopathies, specifically focusing on PERK’s influence on tau protein aggregation. We performed structural modeling and bioinformatic analyses to analyze how tauopathy-associated *PERK* variants impact function. We evaluated how tau aggregation affected PERK signaling in a cell culture model and tested how chemical modulation of the PERK signaling pathway impacted tau aggregation. Last, we analyzed the status of PERK signaling and compared to tau neuropathology in Alzheimer’s disease brains. Our findings support that the PERK pathway prevents tau protein aggregation. Conversely, interfering with PERK pathway signaling increases tau aggregation.

## Results

### Population Distribution of Disease-Associated *PERK* Variants

Genetic studies identify PERK as a disease gene in WRS(26), PSP(13–15), and some forms of Alzheimer’s disease(16,17). To gain insights into PERK’s association with such diverse diseases, we examined the distribution and molecular differences of PERK disease variants in the human population. We identified 1294 variants of human *EIF2AK3/PERK* in gnomAD database (gnomAD v.2.1.1; Genome build: GRCh37 / hg19; Ensembl gene IDENSG00000172071.7) that introduced missense and nonsense changes in coding exons, as well as targeted many non-coding regions (Fig. 1a and Suppl. Data 1). Almost all variants (1270/1294) were ultra-rare with allelic frequencies below 0.1% (Fig. 1b and Suppl. Data 1). Fourteen variants were rare with allelic frequencies between 0.1% to 1% (Fig. 1b and Suppl. Data 1). The remaining 10 *PERK* variants were common with >1% frequency (Fig. 1b and Suppl. Data 1). WRS-associated *PERK* variants all arose at ultra-rare frequencies and introduced nonsense (25 variants) or missense (12 variants) changes exclusively (Fig. 1d and Suppl. Data 1)(27). By contrast, tauopathy-associated *PERK* variants included common, rare, and ultra-rare variants that introduced missense changes or affected non-coding regions but did not cause nonsense changes (Fig. 1e and Suppl. Data 1). Interestingly, the most common *PERK* variant, Haplotype B(14,29), originally identified as a tauopathy risk factor(13), showed striking differences in frequency between racial/ethnic groups (Fig. 1c and Suppl. Data 2), ranging from 5% in African individuals to 49% in East Asian individuals. Conversely, the protective Haplotype A *PERK* variant showed an inverse frequency in these populations (Fig. 1c and Suppl. Data 2).

**Figure 1.**
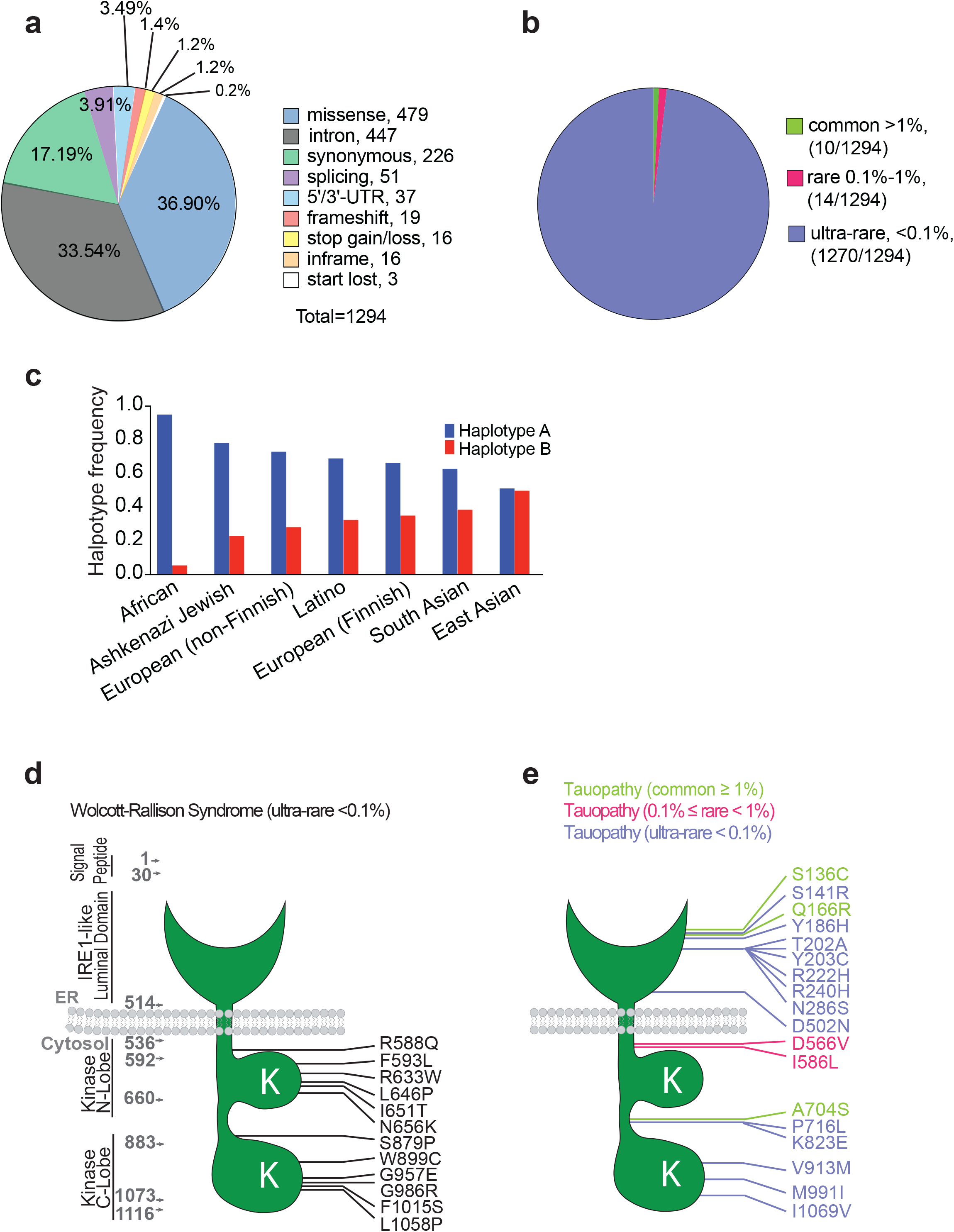
Molecular function and population distribution of PERK variants in human diseases. (**a**) Pie chart shows the molecular functional classification of 1294 genetic human *PERK* variants reported in Genome Aggregation Database (gnomAD). (**b**) Pie chart shows the frequency of human *PERK* variants reported in gnomAD. 1270 *PERK* variants are ultra-rare (<0.1% frequency). 14 *PERK* variants are rare (0.1% to 1% frequency). 10 variants are common (>1% frequency). (**c**) Prevalence of the 2 most common *PERK* variants, *Haplotype A* and *Haplotype B*, across 7 racial/ethnic groups found in gnomAD. The frequency of tauopathy-risk variant, *Haplotype B*, ranges from ∼49% in East Asian populations to ∼5% in African population. Conversely, the frequency of tauopathy-protective variant, *Haplotype A*, ranges from 94% in African population to ∼50% in East Asian populations. (**d, e**) PERK protein cartoons show positions of missense variants linked to WRS (**d**) and tauopathies (**e**) as reported in gnomAD database and NCBI since 2000. Functional domains of the 1116 amino acid human PERK protein include IRE1-like luminal ER stress-sensing domain; ER transmembrane domain; and amino (N) and carboxyl (C) kinase lobes in the cytoplasmic domain. PERK missense variants linked to WRS are all ultra-rare and target PERK’s N- and C-kinase lobes. PERK missense variants linked to tauopathies include common (green), rare (red), and ultra-rare (blue) variants. Tauopathy PERK missense variants frequently map to PERK’s luminal domain; do not overlap with WRS variants; and do not target the kinase lobes.

Next, we focused on the *PERK* variants that introduced missense changes in the protein. The human PERK protein is a 1116-amino acid type 1 integral membrane protein embedded in the endoplasmic reticulum (ER) with a luminal ER stress-sensing domain, coupled to a cytosolic kinase domain(18). The 12 missense variants linked to WRS all targeted the kinase domain (Fig. 1d). By contrast, the 10 tauopathy-associated missense variants targeted the ER stress-sensing luminal domain and less frequently affected cytosolic residues (Fig. 1e). No overlap was found between WRS and Tauopathy-associated PERK variants. These genetic observations suggest that disruption of PERK kinase function underlies the pathogenesis of WRS. By contrast, in tauopathies, kinase function is preserved, but PERK’s ER stress sensing-domain is altered.

### Tauopathy-associated PERK variants disrupt hydrogen bond formation in the ER stress-sensing luminal domain

High resolution mouse and human PERK luminal domain crystal structures(30,31) enable modeling of the impact of tauopathy-risk PERK luminal domain variants on PERK’s ER stress sensing domain. We focused on two luminal domain residues, S136 and R240, because they are well conserved between mammalian PERK proteins (Fig. 2a, 2b). A S136C conversion was present in the Haplotype B PERK tauopathy risk variant(29), and an R240H conversion was found independently as a risk variant in Alzheimer’s disease cohorts(17). Neither of these amino acid substitutions are reported in other mammalian PERK proteins (Fig. 2a, 2b). When we modeled the S136 and R240 residues onto the mammalian PERK luminal domain structure (PDB ID: 4YZY) using PyMol, we observed a direct hydrogen bond (H-bond) between S136 and R240 and 6 additional H-bond interactions formed by surrounding residues, L111, S134, V138, and Q242 (Fig. 2c). Structural modeling predicted that the combination of the C136 risk variant with the R240 protective variant was still able to form 1 direct H-bond but only 4 H-bonds were formed by surrounding residues, S134, G135, and Q242 (Fig. 2d). The combination of the S136 protective variant with the H240 risk variant lost direct H-bond formation but retained 4 H-bonds between L111, S136, V138, and Q242 (Fig. 2e). Last, structural modeling predicted that the combination of a C136 risk variant and H240 risk variant was unable to form direct H-bonds, and only 2 H-bonds could form from surrounding residues, S134 and Q242 (Fig. 2f). In sum, structural modeling of these 2 human variants on the mammalian PERK luminal domain structure revealed a negative impact of disease-associated tauopathy conversions at the 136 and 240 residues upon H-bond formation. The protective variants generated 7 potential H-bonds, but introduction of disease variants impaired H-bond formation between these two residues and the local structural environment. H-bonds stabilize tertiary PERK protein conformation(30). The functional consequences of loss of H-bonds on PERK’s luminal domain ER stress-sensing properties are unclear. However, bioinformatic algorithms predict both conversions to be pathogenic (Fig. 2g). PolyPhen-2, PROVEAN, MutationTaster, SIFT, and CADD found R240H to be pathogenic; and the S136C conversion was pathogenic when analyzed by SIFT and CADD (Fig. 2g). Put together, these analyses support that tauopathy-associated PERK variants negatively impact the structure of the ER stress-sensing luminal domain with predicted pathologic consequences.

**Figure 2.**
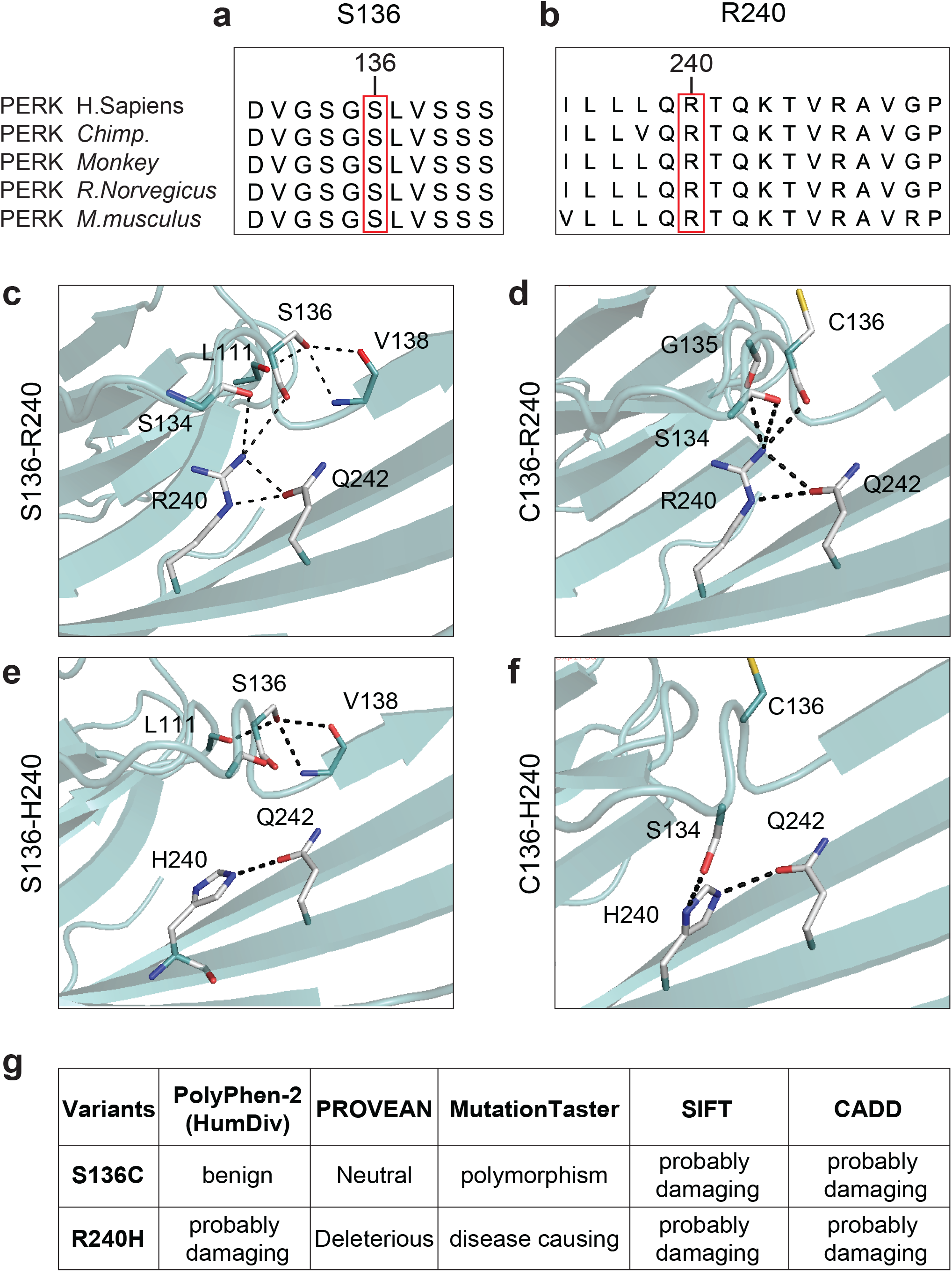
Tauopathy-associated variants disrupt hydrogen bonds in the PERK luminal domain. (**a, b**) Amino acid sequence alignments show conservation of human tauopathy protective PERK variants, S136 and R240, with chimpanzee, monkey, rat, and mouse PERK proteins. Human tauopathy PERK risk variants, C136 and H240, are not reported in other mammalian PERK proteins. (**c, d, e, f**) Hydrogen bonds formed by the PERK 136 and 240 residues were modeled by PyMol using the mouse PERK luminal domain crystal structure (PDB: 4yzy; MMDB ID: 129295). (**c**) The combination of the protective PERK S136 and R240 variants forms 7 H-bonds (dashed lines) locally including 1 direct S136-R240 H-bond. (**d**) The combination of the risk PERK C136 and protective R240 variants forms 5 H-bonds locally including 1 direct C136-R240 H-bond. (**e**) The combination of the protective PERK S136 and risk H240 variants forms 4 H-bonds locally. S136-H240 cannot form direct H-bonds. (**f**) The combination of risk PERK C136 and H240 variants forms 2 H-bonds. C136-H240 cannot form direct H-bonds. White=Carbon; Blue=Nitrogen; Red=Oxygen; Yellow=Sulfur (**g**) Pathogenicity of PERK S136C and R240H missense changes was bioinformatically assessed by 5 algorithms: PolyPhen-2 (HumDiv), PROVEAN, MutationTaster, SIFT, and CADD. PERK S136C was pathogenic using SIFT and CADD. PERK R240H was pathogenic in all algorithms.

### Tau aggregation does not induce ER stress or ERAD and negatively impacts PERK- and IRE1-mediated gene expression in vitro

Tau protein aggregation is a defining feature of tauopathies. We next examined how tau protein aggregation affects PERK signaling and related ER stress-induced processes. We turned to an *in vitro* HEK293 cell model of tau aggregation, “Biosensor” cells that stably express TauRD(P301S)-YFP at low levels diffusely in the cytosol(32–34). When transfected with tauopathy brain protein lysates, the TauRD(P301S)-YFP aggregates into distinct fluorescent puncta(32–34). We prepared brain protein lysates from the PS19 tauopathy mouse model that expresses human P301S tau protein throughout the nervous system(35) and from wild-type mice (Fig. 3a). We confirmed abundant pathologic human tau protein in PS19 mice brain lysates that induced fluorescent tau aggregates when transfected into Biosensor cells (Fig. 3a, 3c, 3d) while wild-type mouse brain lysates showed none of these properties (Fig. 3a, 3c, 3d). We then examined the expression levels of a panel of 31 PERK-regulated genes during tau aggregation in this model using RNA-Seq. These genes were previously shown to be regulated by PERK(36,37), and we also verified robust induction of these genes in thapsigargin-treated Biosensor cells (Suppl. Fig. 1a, 1b; Suppl. Data 3; Suppl Data 4). When we examined PS19 brain lysate-treated Biosensor cells, we saw no induction, but instead, observed a small but significant reduction in expression of the PERK-regulated gene set (Fig. 3f, \*\*\*\**p*≤0.0001, one sample t-test and two-tailed Wilcoxon Signed Rank Test). Examination of 5 individual PERK-regulated genes, *ATF3, RELB, ASNS, GADD34* and *GADD45A*, also showed reduction in expression in PS19 brain-treated cells vs controls (Fig. 3e, \*\**p*≤0.01, two-tailed Student’s *t-*test). Expression of the *PERK* gene itself was not changed in PS19 vs wild-type brain-treated cells (Fig. 3e). Next, we examined the status of IRE1- and ATF6-signaling in Biosensor cells during tau aggregation in our RNA-Seq datasets. For IRE1 signaling, we examined 32 genes previously demonstrated to be regulated by IRE1(36,38). We verified their robust induction in thapsigargin treated Biosensor cells (Suppl. Fig. 1c, 1d; Suppl. Data 3; Suppl. Data 4). When we examined PS19 brain lysate-treated cells, we also saw a significant reduction in expression of the IRE1-regulated gene set (Fig. 3h, \*\*\*\**p*≤0.0001, one sample t-test and two-tailed Wilcoxon Signed Rank Test). Examination of selected individual IRE1-target genes, *ERdj4, SLC3A2, VEGFA, UFM1*, confirmed significant reduction in gene expression (Fig. 3g, \*\**p*≤0.01, two-tailed Student’s *t-*test). *IRE1* gene expression itself was not changed (Fig. 3g). For ATF6 signaling, we examined expression of 74 genes previously reported to be induced by ATF6(36,39,40) and also robustly upregulated in Biosensor cells after thapsigargin treatment (Suppl. Fig. 1e, 1f; Suppl. Data 3; Suppl Data 4). By contrast to the PERK- and IRE1-gene panels, the ATF6-regulated genes showed no significant changes with PS19 brain treatment (Fig. 3i, 3j). Last, we analyzed a 74 gene ER stress-associated degradation (ERAD) panel present in the GO database ERAD term (GO:0036503) (http://amigo.geneontology.org/amigo/term/GO:0036503). As expected, we found significant induction of the ERAD gene set in Biosensor cells after thapsigargin treatment (Suppl. Fig. 1g). By contrast, ERAD gene expression was not altered with PS19 brain treatment (Fig. 3k, 3l). Bioinformatic pathway analysis on the entire RNA-Seq dataset using gene ontology (GO) and gene set enrichment analysis (GSEA) further confirmed significant induction of ER stress and ERAD in thapsigargin-treated Biosensor cells (Suppl. Fig. 1h; Suppl. Fig. 2); while PS19 brain treatment did not induce ERAD, and ER stress was not changed by GSEA and showed significant down-regulation by GO analysis (Fig. 3l; Suppl. Fig. 2). Consistent with these transcriptomic findings, we found pronounced increase in phosphorylated eIF2*α*levels in thapsigargin-treated Biosensor cells that was not seen in PS19 brain-treated lysates (Suppl. Fig. 1i). In sum, these results demonstrate that tau aggregation does not trigger ER stress or ERAD in the Biosensor cell culture model. Instead, tau aggregation is associated with a small but significant down-regulation of PERK-mediated gene expression.

**Figure 3.**
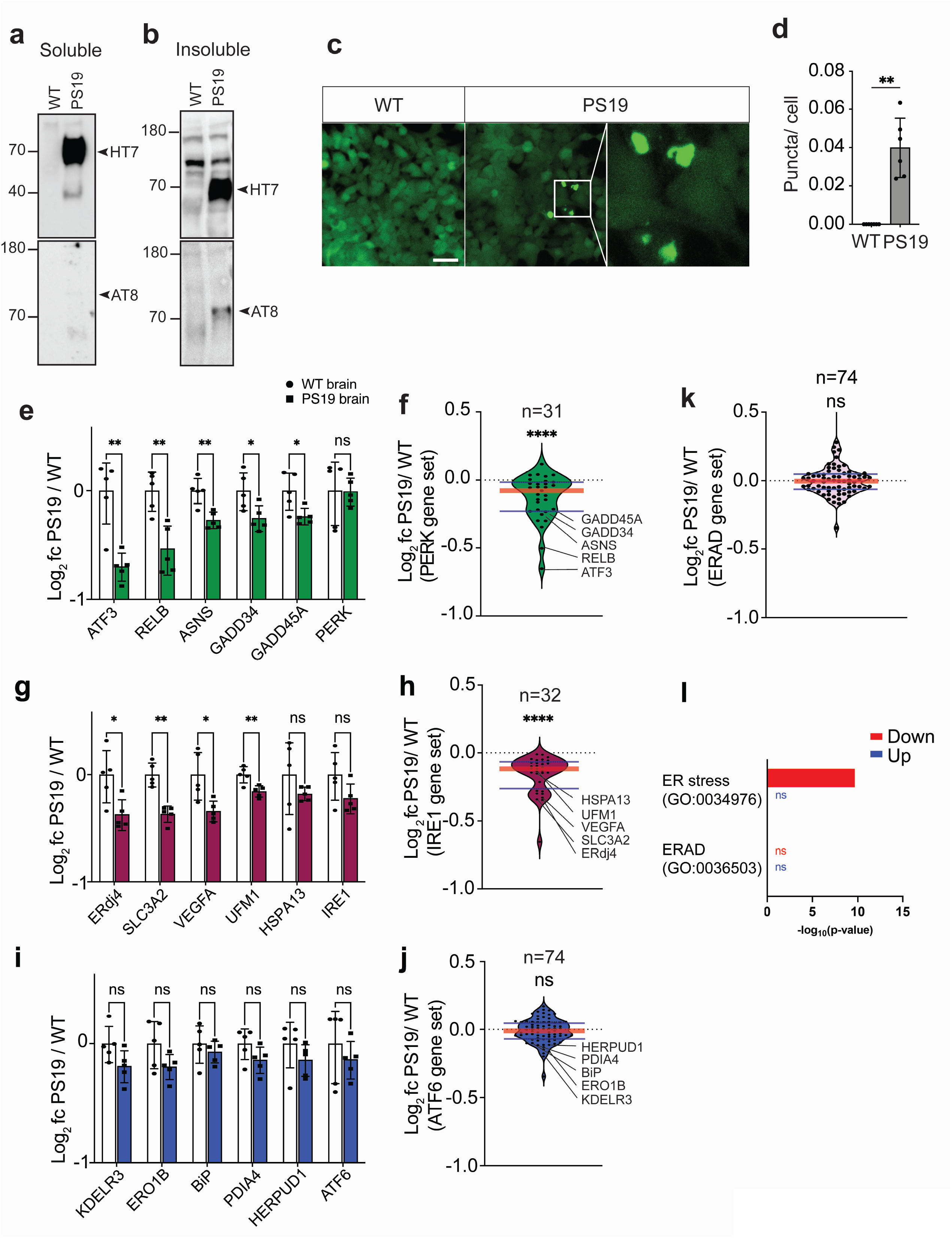
Tau aggregation in cell culture does not induce ER stress. (**a, b**) Protein lysates were prepared from wild-type and PS19 mouse brains. Soluble and insoluble protein lysate fractions were immunoblotted for total human Tau (HT7) and phospho-human Tau (AT8). Arrowheads mark positions of tau protein. (**c**) Biosensor cells were transfected with wild-type or PS19 soluble brain lysate. After 24 hours, Tau-YFP aggregates (puncta) were imaged by fluorescent microscopy. The white box outlines magnified image of Biosensor cells with puncta. Scale bar = 50 μm. (**d**) Fluorescent puncta were quantified after wild-type or PS19 mouse brain lysate transfection, and the puncta number was normalized to cell number (\*\**p*≤0.01, one-tailed Student’s *t-*test, n=6 independent transfections, Mean ± S.D.). (**e-l**) The mRNA levels of PERK-, IRE1-, ATF6-, and ERAD-regulated genes were examined by RNA-Seq of Biosensor cells transfected with wild-type or PS19 mouse brain protein lysate for 24h. (**e, f**) Gene expression levels of 31 PERK-regulated genes in PS19 brain lysate-treated cells relative to wild-type brain lysate-treated are shown as log_2_ fold change. The graph (**e**) shows levels of the 5 PERK-regulated genes most significantly reduced between wild-type and PS19-treated cells and expression levels of the *PERK* gene itself. The violin plot (**f**) shows levels of the entire PERK-regulated gene set. (**g, h**) Gene expression changes of 32 IRE1-regulated genes in PS19-treated cells relative to wild-type treated are shown. The graph (**g**) shows levels of the 5 IRE1-regulated genes most differentially expressed between wild-type and PS19-treated cells and expression levels of *IRE1* gene itself. The violin plot (**h**) shows levels of the entire IRE1-regulated gene set. (**i, j**) Gene expression changes of 74 ATF6-regulated genes in PS19 brain lysate-treated cells relative to wild-type treated are shown. The graph (**i**) shows levels of the 5 ATF6-regulated genes most differentially expressed between wild-type and PS19-treated cells and expression levels of ATF6 gene itself. The violin plot (**j**) shows levels of the entire ATF6-regulated gene set. (**k**) The violin plot shows levels of 74 ERAD-regulated genes. (**l**) GO analysis identifies significantly decreased ER stress term (GO:0034976) in PS19-treated Biosensor cells. ERAD pathway (GO:0036593) shows no significant (ns) change after PS19 brain treatment. Error bars in **e**, **g**, and **i** represent mean ± standard deviation. Black circles and squares represent 5 independent experimental replicates. (* *p*≤0.05, ** *p*≤0.01, not significant (ns), two-tailed Student’s *t-*test). The red horizontal line in figures **f**, **h**, **j, k** marks the median level of expression of the gene set, and the thin horizontal blue lines delimit upper and lower gene expression quartiles in the violin plots. (\*\*\*\**p*≤0.0001, one sample t-test and two-tailed Wilcoxon Signed Rank Test, n=5 experimental replicates). Detailed gene expression RNA-Seq information is shown in Supplementary data 3.

### Genetic and chemical inhibition of PERK pathway promotes Tau aggregation

Our genetic RNA-Seq experiments in Biosensor cells identified a correlation between tau aggregation and reduced PERK signaling. To test for causality between PERK pathway signaling and tau aggregation, we added small molecules that target PERK, GADD34, or eIF2B to Biosensor cells undergoing tau aggregation. We used PERK kinase inhibitors, GSK2656157, GSK2606414, or ISRIB, which locks eIF2B in an active state (41–43). To increase p-eIF2a levels, we added GADD34 inhibitor, salubrinal(44), to PS19-brain transfected cell media. We found significant increase in tau fluorescent puncta with addition of GSK2656157, GSK2606414, or ISRIB, while salubrinal treatment significantly reduced the formation of fluorescent puncta (Fig. 4a, 4b). Furthermore, we found that addition of salubrinal to PERK inhibitor GSK2606414-treated cells could antagonize the formation of the fluorescent tau aggregates (Fig. 4c, 4d). These studies show that inhibition of PERK kinase or locking eIF2B in an active state directly promotes tau aggregation, while increased eIF2α phosphorylation prevents tau aggregation in Biosensor cells.

**Figure 4.**
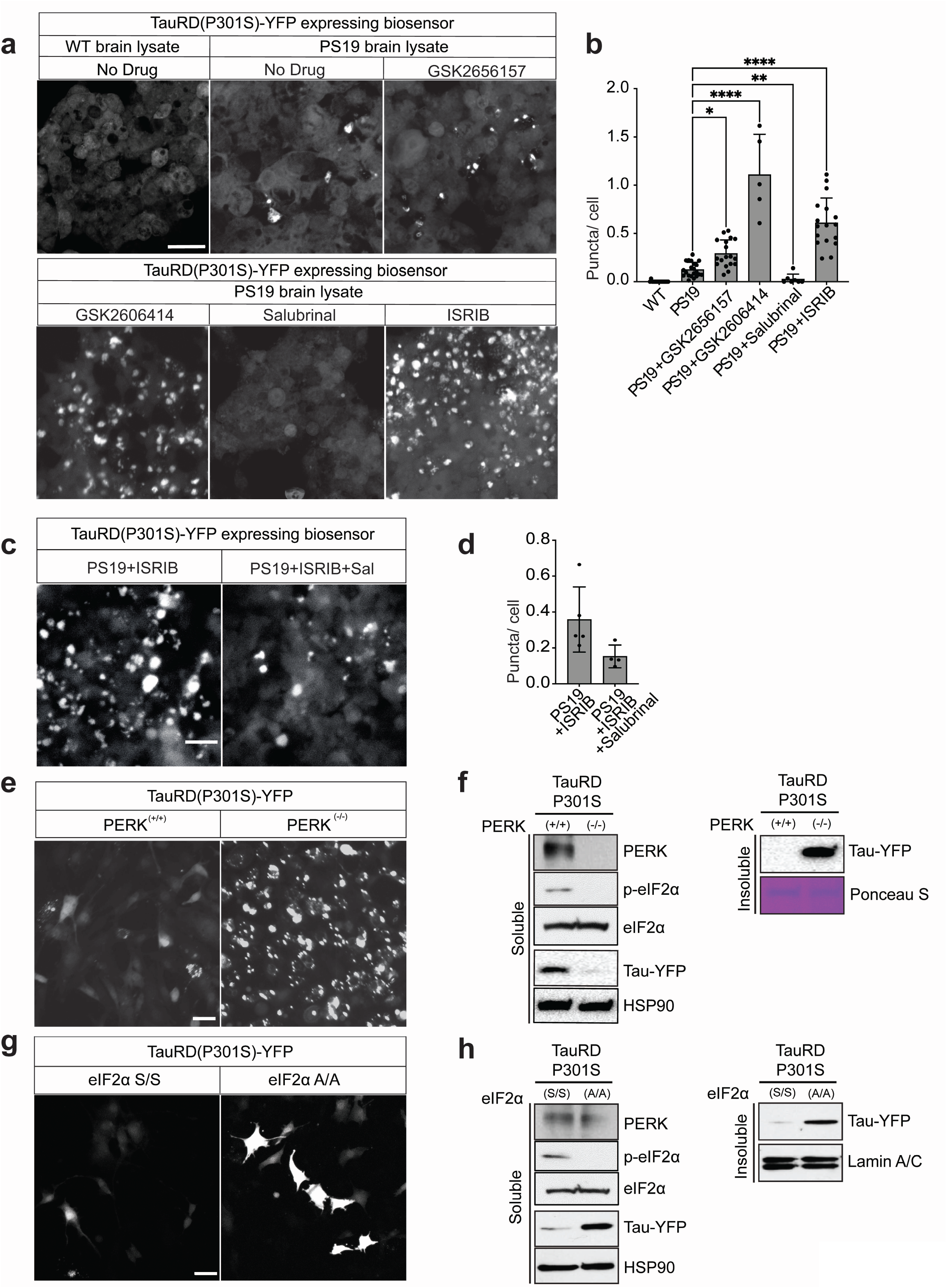
PERK signaling prevents Tau-YFP aggregation. (**a**) Biosensor cells were transfected with wild-type or PS19 brain lysate and co-incubated with GSK2656157 (5uM), GSK2606414 (5uM), Salubrinal (2.5uM), or ISRIB (5uM). After 24 hours, Tau-YFP fluorescent puncta were imaged by fluorescent microscopy. Scale bar = 30 μm. (**b**) Quantification of puncta number from (**a**) normalized by cell number. *p*-value was calculated by two-way ANOVA Tukey’s multiple comparisons test, Mean± S.D. * *p*≤0.05, ** *p*≤0.01, \*\*\*\**p*≤0.0001 (n ≥ 5 experimental replicates). (**c**) Biosensor cells were transfected with PS19 brain lysate and co-incubated with ISRIB for 24 hours. Media was replaced ± Salubrinal for another 24 hours. Tau-YFP fluorescent puncta were imaged by microscopy after these drug treatments. scale bar = 25μm. (**d**) Quantification of puncta number from (**c**) normalized by cell number. (**e, f**) *PERK^-/-^* or *PERK^+/+^* MEFs were transduced with TauRD(P301S)-YFP. After 48 hours, cells were imaged by fluorescence microscopy (**e**, scale bar = 25μm), and protein lysates were prepared. (**f**) Soluble protein fractions were immunoblotted for Tau-YFP and HSP90 (loading control). Insoluble fractions were immunoblotted for Tau-YFP and Ponceau stained (loading control). (**g, h**) *eIF2α^A/A^* or *eIF2α^S/S^* MEFs were transduced with TauRD(P301S)-YFP. After 48 hours, cells were imaged by fluorescence microscopy (**g**, scale bar = 25μm), and protein lysates were prepared. (**h**) Soluble protein fractions were immunoblotted for Tau-YFP or HSP90 (loading control). Insoluble fractions were immunoblotted for Tau-YFP and Lamin A/C (loading control).

To further test the relationship between PERK signaling and tau aggregation, we transduced TauRD(P301S)-YFP into *PERK^+/+^* and *PERK^-/-^* mouse embryonic fibroblasts (MEFs)(45). In *PERK^+/+^* MEFs, the fluorescence signal from the TauRD(P301S)-YFP construct was diffusely found in cell soma, while in *PERK^-/-^* MEFs, the fluorescent signal was found in intense puncta (Fig. 4e). When we examined protein lysates from these transduced *PERK* MEFs, we saw that TauRD(P301S)-YFP protein redistributed from the soluble fraction in *PERK^+/+^* MEFs to insoluble fraction in *PERK^-/-^* MEFs (Fig. 4f). Next, we transduced TauRD(P301S)-YFP into *eIF2α^S/S^* (wild-type) and *eIF2α^A/A^* (ablation of phosphorylation on 51 amino acid) MEFs(46). Similar to results seen in *PERK^+/+^* and *PERK^-/-^* MEFs, TauRD(P301S)- YFP fluorescence intensity was dramatically increased in *eIF2α^A/A^* MEFs as compared to *eIF2α^S/S^* MEFs (Fig. 4g) although puncta were less visible. Protein lysates from transduced *eIF2α^S/S^* and *eIF2α^A/A^* cells showed that TauRD(P301S)-YFP protein was increased in both soluble and insoluble fractions (Fig. 4h). Immunoblot analysis of eIF2*α* phosphorylation and PERK confirmed absence of both proteins in *PERK^-/-^* and absence of p-eIF2*α* in *eIF2α^A/A^* MEFs (Fig. 4f, 4h). Taken together, these results support that increased eIF2 phosphorylation prevents tau aggregation in vitro, and that impaired PERK kinase activity or loss of eIF2 phosphorylation increases tau aggregation.

Last, we tested how chemical modulation of other UPR pathways influenced tau aggregation. When we treated PS19-transfected Biosensor cells with IRE1 inhibitor, 4u8c(47), or ATF6 inhibitor, Ceapin-A7(48), we saw significant increase in fluorescent tau puncta compared to controls (Fig. 5a, 5b). By contrast, addition of an ATF6 pathway activator, AA147(49) did not significantly alter the number of fluorescent tau puncta in PS19 brain lysate-transfected cells (Fig. 5a, 5b). Together, these findings support that inhibition of IRE1 and ATF6, like inhibition of PERK signaling, also increases tau aggregation *in vitro*.

**Figure 5.**
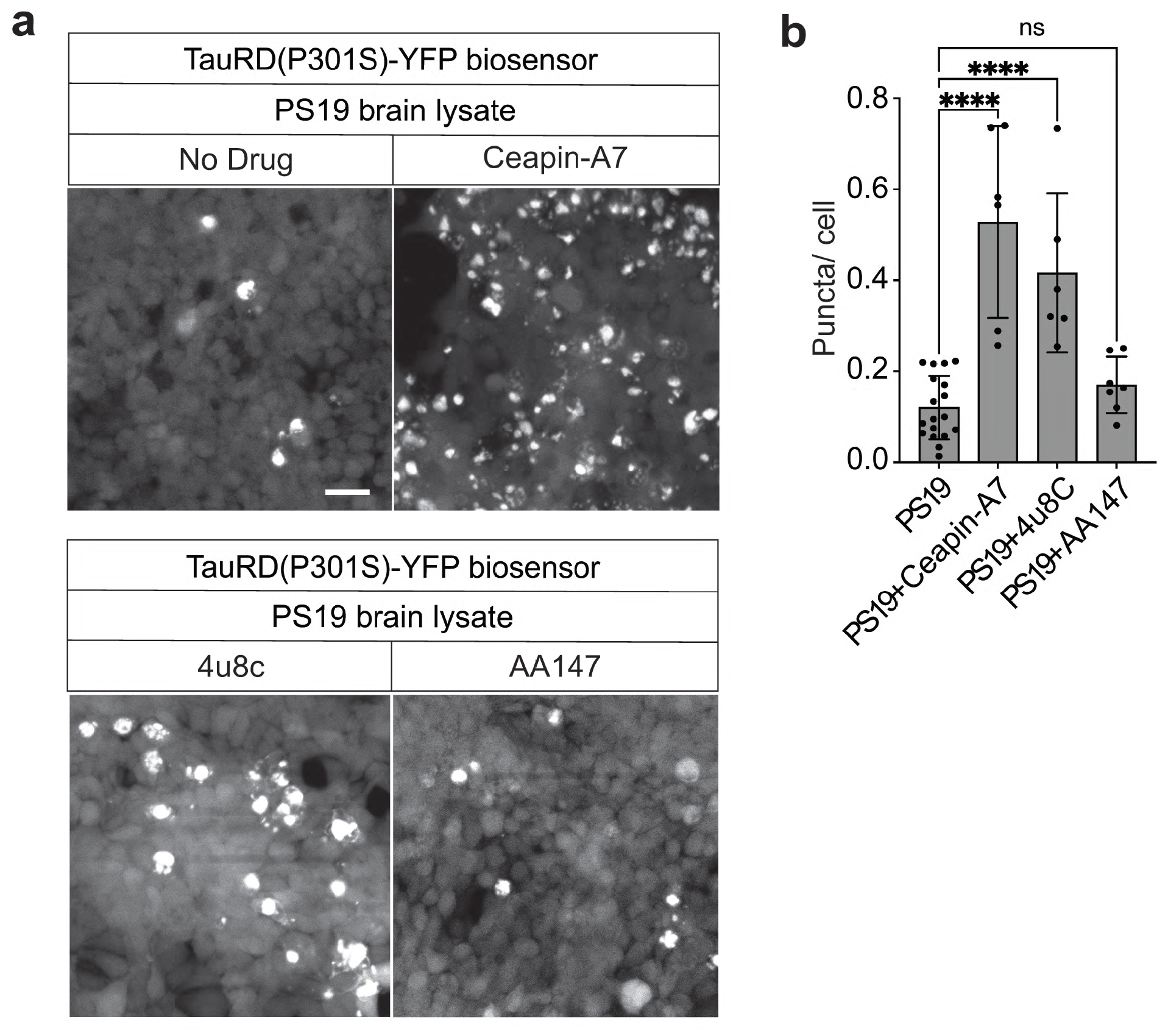
IRE1 and ATF6 pathway inhibitors cause Tau-YFP aggregation in cell culture. Biosensor cells were transfected with PS19 brain lysate and co-incubated with ATF6 inhibitor Ceapin-A7 (10uM), IRE1 inhibitor, 4u8c (10uM), or ATF6 pathway activator, AA147 (10uM). After 24 hours, Tau-YFP fluorescent puncta were imaged by fluorescent microscopy. Scale bar = 30μm. (**b**) Quantification of puncta number from (**a**) was normalized by cell number. *p*-value was calculated by two-way ANOVA Tukey’s multiple comparisons test, Mean± S.D., not significant (ns), \*\*\*\**p*≤0.0001 (n ≥ 5 experimental replicates).

### PERK signaling is reduced in Alzheimer’s Disease patient hippocampi

Our *in vitro* studies support that reduced PERK signaling leads to increased tau aggregation. To investigate if this relationship between PERK activity and tau aggregation was found *in vivo*, we biochemically analyzed PERK signaling status in frozen hippocampi banked from postmortem brain donors with Braak-staged tau neuropathology(50,51). We obtained 5 hippocampi samples from patients with clinical history of Alzheimer’s disease and Braak VI tau neuropathology (Table 1) and compared with 5 Braak I control hippocampi (patients with no dementia and no tau neuropathology) (Table 1) We prepared soluble and insoluble protein fractions(52) and confirmed increased pathologic tau protein in Braak VI compared to Braak I samples (Fig. 6a, 6b). We further confirmed that Braak VI, but not Braak I, brain lysates induced fluorescent puncta formation (tau aggregation) in Biosensor cells (Fig. 6c) consistent with prior studies(32-34,53,54). Next, we examined PERK protein expression in these Braak I and VI samples. In the soluble fractions, we found statistically significant reduction of phospho-PERK levels in Braak VI compared to Braak I samples while total PERK levels did not differ (Fig. 6d, 6e, 6f). PERK phosphorylation is a marker of PERK activation(18), and this finding in Braak I and Braak VI brains provides *in vivo* evidence that reduced PERK pathway activity correlates with increased tau aggregation.

**Figure 6.**
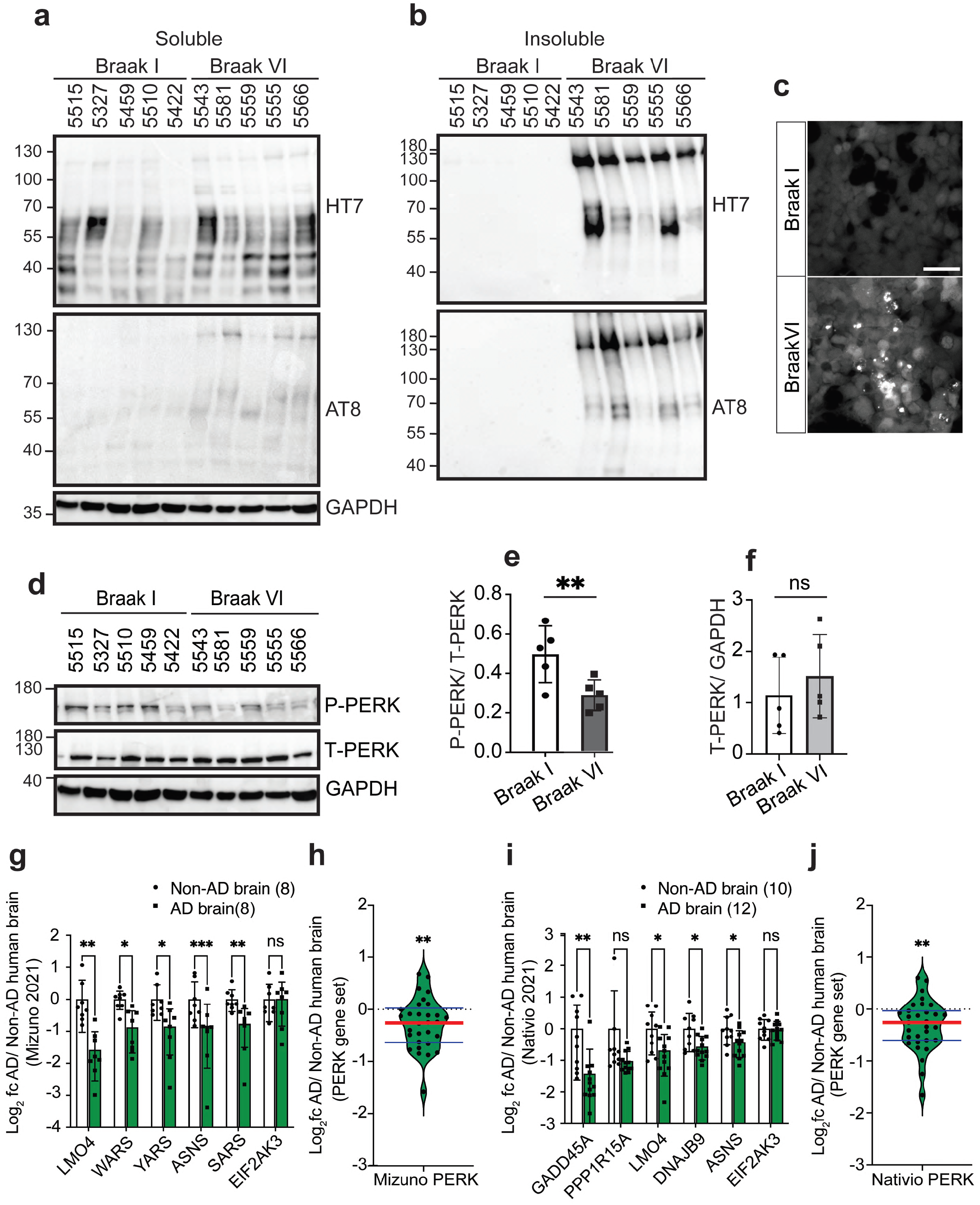
PERK pathway activity is reduced in Alzheimer’s disease patient brains. (**a, b**) Protein lysates were prepared from 1mg of 5 Braak I (normal) and 5 Braak VI (Alzheimer’s disease) patient hippocampi. Soluble (**a**) and insoluble (**b**) protein lysate fractions were immunoblotted for total human Tau (HT7), phospho-Tau (AT8), and GAPDH (loading control). Individual brain identification numbers are listed above the blots, and associated clinicopathology information are in Table 1. (**c**) Biosensor cells were transfected with normal (Braak I) or Alzheimer’s disease (Braak VI) soluble brain lysate. After 24 hours, Tau-YFP aggregates (fluorescent puncta) were imaged by fluorescent microscopy. Scale bar = 50μm. (**d, e, f**) Soluble brain lysates from (**a**) were immunoblotted for phospho-PERK, and total PERK; protein levels were quantified by densitometry and normalized by loading controls, PERK (d) and GAPDH (a). Phospho-PERK and total PERK were not detected in insoluble fraction (data not shown). \*\**p* ≤ 0.01, not significant (ns), two-tailed Student’s *t-*test. Mean ± S.D. (**g, h**) Gene expression levels of PERK-regulated genes in hippocampi of AD brains (n=8) relative to non-AD brains (n=10) from GSE173955 (2021 Mizuno) are shown as log_2_ fold change. The graph (**g**) shows levels of the 5 PERK-regulated genes most significantly reduced between AD brains and non-AD brains, and expression levels of *PERK* gene itself. The violin plot (**h**) shows levels of the entire PERK-regulated gene set. Detailed clinicopathology information are available from (2021 Mizuno) and summarized in Table 1. (**i, j**) Gene expression levels of PERK-regulated genes from hippocampi of AD brains (n=12) relative to non-AD brains (n=10) from 2020 Nativio (GSE159699) are shown as log_2_ fold change. The graph (**i**) shows levels of the 5 PERK-regulated genes most significantly reduced between AD and non-AD brains and *PERK* gene expression levels itself. The violin plot (**j**) shows levels of the entire PERK-regulated gene set. Detailed clinicopathology information of these AD and non-AD brains are available at 2020 Nativio and summarized in Table 1. Error bars in **g**, and **i** represent mean ± standard deviation. Black circles and squares represent individual AD or non-AD brains. (* *p*≤0.05, ** *p*≤0.01, *** *p*≤ 0.001, not significant (ns), two-tailed Student’s *t-*test). The red horizontal line in figures **h**, **j** marks the median level of gene expression, and the thin horizontal blue lines delimit upper and lower gene expression quartiles in the violin plots. (\*\**p*≤0.01, one sample t-test and two-tailed Wilcoxon Signed Rank Test).

To further investigate if PERK signaling was reduced in Alzheimer’s disease patient brains, we analyzed mRNA levels of the PERK-regulated gene set in RNA-Seq datasets collected from Alzheimer’s disease hippocampi and normal hippocampi (Fig. 6g, 6h, 6i, 6j). We were not able to perform RNA-Seq on the same brain cases used for immunoblotting, because the protein preparation methods were not compatible with the preservation of high-quality RNA. Instead, we examined hippocampal RNA-Seq datasets independently generated from two Alzheimer’s disease patient cohorts in Japan and the USA(55,56). The first RNA-Seq dataset contained 8 clinically and neuropathologically Braak-staged Alzheimer’s brain hippocampi matched with 8 control hippocampi collected from Japanese patients ((56) and Table 1). The second RNA-Seq dataset contained 12 clinically and neuropathologically Braak-staged Alzheimer’s brain hippocampi and 10 normal control hippocampi collected from an American cohort ((55) and Table 1). When we examined the expression levels of the PERK-regulated gene set in these two hippocampal RNA-Seq datasets, we found significantly reduced levels of the PERK-regulated gene set as well as individual PERK-regulated genes in Alzheimer’s disease brains compared to controls in both Japanese (Fig. 6g, 6h, Suppl. Data 5) and American cohorts (Fig 6i, 6j, Suppl. Data 5). Together, our biochemical studies and analyses of published RNA-Seq datasets from Alzheimer’s hippocampi provide *in vivo* support that PERK pathway activity inversely correlates with tau pathology.

## Discussion

PERK is a key effector of the UPR and ISR, and controls translational and transcriptional programs that impact vital cellular processes including amino acid metabolism, antioxidative response, ER protein folding, autophagy, and apoptosis. In people, ultra-rare *PERK* variants that ablate PERK function are causally linked to the autosomal recessive disease WRS. Another distinct group of more common and non-overlapping *PERK* variants increases risk for tauopathies. Previously, we found that tauopathy-associated *PERK* variants diminished PERK signaling *in vitro*(28), but the consequences of changes in PERK pathway activity on tauopathy pathogenesis remained unclear. Here, we found that impaired PERK pathway activity increased tau aggregation, while increasing p-eIF2a reduced tau aggregation in tauopathy cell culture models. We found that impaired PERK pathway activity correlated with increased tau neuropathology in brain tissues from three different tauopathy patient cohorts. We found that tauopathy-associated PERK variants did not target kinase activity, but instead, at least two variants negatively impacted the tertiary structure of the ER stress-sensing luminal domain.

Based on these findings, we propose that tauopathy-associated PERK variants increase disease risk, in part, by facilitating tau protein aggregation and downstream neuropathology. Our findings showing causality between PERK signaling and tau aggregation were based on experiments performed in MEFs ablated for PERK or eIF2*α* phosphorylation or in HEK293 cells expressing fluorescent protein-tagged TauRD(P301S). These are robust and reproducible systems to study tau protein aggregation(32-34,53,54), but the small TauRD(P301S) fragment likely does not recapitulate all aspects of aggregation by native full-length tau protein isomers. A related limitation of our study is that the MEF and HEK293 cell models do not recapitulate the neuronal and glial environments where tau causes neuropathology. It will be important to evaluate how tauopathy-associated PERK variants impact tau pathology in native neural cell types. Despite the limitations of our abbreviated tau construct and cell culture models, our analyses of PERK signaling in primary patient brain tissues support that reduced PERK pathway activity correlates with increased tau protein aggregation and pathology. Given the absence of therapies for tauopathies and the potential of small molecule PERK and ISR pathway agents to influence tau aggregation, the role of PERK signaling warrants further investigation in the pathogenesis and treatment of tauopathies.

An unexpected finding in our analysis of human *PERK* variants was the unequal distribution of the tauopathy-risk *Haplotype B* allele between racial/ethnic groups ranging from a low of ∼5% in African populations up to 49% in East Asian populations (Fig. 1c). Prior studies found increased prevalence of PSP in a Japanese cohort (17.90 per 100,000 people)(57), compared to 6 per 100,000 in a European cohort(58,59); and 2.95 per 100,000 in the US population. We speculate that the increased prevalence of PSP in the Japanese populace could reflect the higher prevalence of the tauopathy disease allele, *Haplotype B*, in this East Asian population. PSP prevalence has not been examined in African groups, but we predict that PSP prevalence should be significantly lower because of the relative rarity of the *Haplotype B* in this population (Fig. 1c). Additional molecular epidemiologic studies across different ethnic/racial groups can shed light upon the link between *Haplotype B* prevalence and distribution of disease.

We observed that many tauopathy-associated PERK variants target the ER stress-sensing luminal domain (Fig 1e), and modeling of amino acid substitutions at residues 136 and 240 on the mammalian crystal structure of the PERK luminal domain revealed disruption of H-bonds between these two residues when converted to disease variants (Fig. 2). How do these changes in the luminal domain impact PERK signaling? The local sequences bearing residues 136 and 240 are important for PERK tetramerization/oligomerization in response to ER stress(30,31), and therefore, we speculate that the S136C and R240H conversions may alter PERK’s tetramerization/oligomerization ability. PERK’s luminal domain also binds to chaperones, and luminal domain PERK variants may also alter interactions with PERK regulatory co-factors like Grp78/BiP(30,60). PERK dimerization after ER stress is less likely to be directly affected by changes at S136 and R240 because these residues are not located near the PERK dimerization interface(30). Based on these modeling observations, we propose that tauopathy variants in the PERK luminal domain interfere with PERK’s ability to accurately sense and respond to ER stress; dysregulation of kinase activation and downstream signaling arises as a secondary consequence.

The status of PERK signaling and ER stress in tauopathy pathogenesis is mixed. Biochemical analysis of tauopathy mouse brains at many ages prior to and during disease revealed no increase of PERK signaling or induction of ER stress in these models(61,62). By contrast, increased PERK signaling was reported in some diseased neurons through immunostaining of primary Alzheimer’s disease and PSP patient brain sections(63,64). In our studies, we saw no activation of PERK signaling in the cell culture tau aggregation model or in primary tauopathy patient hippocampi (Fig. 3, Fig. 6). Our PERK signaling analysis was performed by Western blots or RNA-Seq of bulk lysates from *in vitro* cultured cells or human brain tissues. This approach may mask increased PERK signaling in individual or small populations of cells. Single-cell approaches can provide better resolution of PERK signaling dysregulation during tau aggregation in distinct neural cell types.

In our studies, we not only saw no activation of PERK signaling but instead, we saw reduced PERK signaling in the HEK293 cell culture model and in tauopathy patient brain samples by gene expression measurements (Fig. 3, Fig. 6). To our knowledge, reduced PERK signaling in tauopathy has not been reported previously. We do not know how or why PERK signaling as determined by transcriptional output is suppressed in the cell culture model of tau aggregation or in advanced tauopathy patient brains. However, impaired PERK function worsens ER homeostasis, increases oxidative stress, and increases protein misfolding(45). In the brain, increased tau aggregation could be a specific deleterious consequence of PERK dysfunction.

Our findings support prior studies that pharmacologic PERK activation or PERK overexpression attenuate tau pathology in vitro and in vivo (65). Augmented PERK signaling, and more broadly UPR/ISR signaling, may provide tools to ensure tau protein homeostasis and prevent the emergence of pathologic tau aggregates. A PERK augmentation strategy would be especially applicable for carriers of tauopathy-associated *PERK* hypomorph alleles.

## Experimental procedures

### Genome data collection and interpretation of PERK variants

1294 PERK variants were examined in three publicly available databases (access date: 05 January 2022): the Genome Aggregation Database (gnomAD version 2.1.1), ClinVar (NCBI), and European Bioinformatics Institute (EMBL-EBI) Database. The 1294 variants included 479 missense variants; 447 intron variants; 226 silent variants; 51 splicing variants; 37 UTR variant; 19 frameshift variant; 19 start and stop variants; 16 in frame variants. 12 WRS and 18 Tauopathy missense mutations were identified in the genome databases and in various publications. https://uswest.ensembl.org/Homo_sapiens/Gene/Phenotype?db=core;g=ENSG00000172071;r=2:88556741-88627464; https://www.ncbi.nlm.nih.gov/CBBresearch/Lu/Demo/LitVar/index.html#!?query=EIF2AK3; https://gnomad.broadinstitute.org/gene/ENSG00000172071?dataset=gnomad_r2_1) (Fig. 1). PERK haplotype race/ethnic frequency was calculated by population allele count divided by allele number from the gnomAD database. Tauopathy related mutations were grouped as common *≥*1%, 0.1%*≤* rare<1%, and ultra-rare<0.1% based on the population allele frequencies identified from gnomAD. Pathogenicity of PERK variants was assessed by publicly accessible Web server-based prediction tools (PolyPhen-2: http://genetics.bwh.harvard.edu/pph2/; PROVEAN: http://provean.jcvi.org/index.php; MutationTaster: https://www.mutationtaster.org/; SIFT: https://sift.bii.a-star.edu.sg/www/SIFT_seq_submit2.html; https://cadd.gs.washington.edu/. All last accessed June 2021). PolyPhen-2 (Polymorphism Phenotyping v2) uses sequence alignments, phylogenetics, and structural data to characterize amino acid substitutions and calculates a score for the variant, classifying it as ‘benign’, ‘possibly damaging’, or ‘probably damaging’. Scores range from 0.0 (benign) to 1.0 (probably damaging). SIFT (sorting intolerant from tolerant) predicts the impact of an amino acid change on protein function by comparing amino acid alignments from related sequences to calculate a ‘SIFT score’: 0–0.05 will be classified as ‘damaging’, 0.05–1 as ‘tolerated’. Prediction of pathological mutations on proteins uses sequence information for its neural network and predicts the effect of amino acid changes on protein function, by calculating a reliability index ranging from 0 to 10 (most unreliable to most reliable prediction) and a prediction of either ‘neutral’ or ‘pathological’. Mutation prediction is based on SIFT and structural and functional properties of proteins. Mutation prediction was created using disease-associated mutations from HGMD and neutral amino acid substitutions from Swiss-Prot. The output contains a general score, (g) where g>0.5 (P<0.05) is actionable, g>0.75 (P<0.05) is confident and g>0.75 (P<0.01) is very confident that an amino acid substitution is likely to have a phenotypic effect. All data collection, pathogenicity assessments, and database annotation were performed by scientists trained with standardized training modules and annual proficiency testing.

### PERK luminal domain structural modeling analysis

Protein sequence alignment was evaluated using the EMBL-EBI EMBOSS Needle program (https://www.ebi.ac.uk/Tools/psa/emboss_needle/) and visualized using JalView (version 2.11.1.4; www.jalview.org). The visualization of the reconstructed model structure and H-bond prediction of the PERK protein was prepared using PyMol molecular graphics system (http://www.pymol.org). The crystal structure of mammalian PERK luminal domain (PDB ID: 4YZY) was used for the structure modeling analysis. When superimposed 4YZY with the mutation models, all H-bonds of residues R240/H240 and S136/C136 were highlighted and shown in dash lines.

### Antibodies and chemicals

Antibodies including HT-7 (#MN1000, Invitrogen), AT8 (# MN1020, Invitrogen), YFP (#ab6556, Abcam), HSP90 (#ab13492, Abcam), Lamin A/C (#2032, Cell Signaling Technology), GAPDH (#ab8245, Abcam), T-PERK (#3192, Cell Signaling Technology), P-PERK (#3179, Cell Signaling Technology), eIF2*α* (# 5324, Cell Signaling Technology), P-eIF2*α* (# 5324, Cell Signaling Technology), were pre-tested to detect the targeted proteins. Small molecules included GSK2656157 (# 9466-5, BioVision); GSK2606414 (#S7307, Selleckchem); Salubrinal (#S2923, Selleckchem); ISRIB (#S7400, Selleckchem); Ceapin-A7 (#SML2330, Sigma-Aldrich); 4u8C (CAS14003-96-4, Calbiochem), and AA147 (Product No. 6538059, ChemBridge) were prepared in DMSO following manufacturer’s instruction and stored at −80°C as stock solution. ER stress-inducing chemical, thapsigargin, was dissolved in DMSO and added to the cell culture media at a concentration of 300 nM. The working solutions were freshly prepared with - 80°C stock solution.

### Mouse and human brain protein lysate preparation

#### Mouse brain extraction

P301S tau transgenic (PS19) mice (B6; C3-Tg(Prnp-MAPT*P301S)PS19Vle/J; stock number: 008169; Jackson Laboratory) harboring T34 isoform of microtubule-associated protein tau with one N-terminal insert and four microtubule binding repeats (1N4R) encoding the human P301S mutation were obtained from Jackson Laboratory and maintained in a C57Bl/6j genetic background in standard vivarium environment (12h light:12h dark cycle). Approved laboratory personnel checked mice during the light phase of the light:dark cycle to determine birthdates and weaned the new pups in 3 weeks. At 6 months, PS19 and WT male littermates were anesthetized with ketamine/xylazine (1mg/kg) followed by intracardiac perfusion with saline. Then, mice were euthanized by carbon dioxide and cervical dislocation, and brains were subsequently removed following institutional guidelines and with IACUC approval. Fresh brains were then homogenized with Dounce homogenizer in ice cold 1X RIPA buffer with protease inhibitor (#S8820, Sigma-Aldrich). Following centrifugation (13,000 rpm) at 4°C for 20min, the soluble fraction was analyzed for protein concentration and frozen. The supernatants were aliquoted and stored at −80°C until further use. The insoluble fraction was vigorously resuspended by vortex and boiled with 4X SDS Sample buffer for 10min. 10μg of total protein was run on an 1-15% Bis/Tris gel and transferred to nitrocellulose for western blotting.

#### Human brain extraction

Frozen human brain tissues were obtained from UC San Diego Alzheimer Disease Research Center (ADRC). The diagnoses and demographics in Table 1 were obtained from patients neurologically and psychometrically studied at the UCSD-ADRC with institutional IRB approval. Patients gave informed consent for postmortem brain sample collection for research purposes. Upon autopsy, patient brains were collected by the UCSD ADRC Neuropathology Core and sagittally divided; the left hemibrain was fixed in 10% buffered formalin for neuropathological analysis for Braak tau staging; and the right hemibrain sections were frozen at −70 °C for subsequent protein isolation. For this study, frozen hippocampal tissues were obtained. Human brain protein lysate extraction followed the previous literature (52). In brief, 0.3g of brain tissue was homogenized on ice with 5.3x volume (w/v) Goedert buffer composed of 10mM Tris-HCl, pH = 7.4, 0.8M NaCl, 1mM EGTA, and 10% sucrose including protease inhibitor, *cOmplete* mini (11836153001, Roche), per 10mL and the remaining solution volume with pure PSMF. The homogenate mixture was then spun at 20,000xg for 20 minutes. The supernatants were retained afterwards and re-homogenized in 2.65x volume (w/v) of Goedert buffer. The homogenates were spun at 20,000xg for 20 minutes. The previous supernatant was then combined with newly retained supernatant, and 20uL supernatant was saved for protein concentration. Afterwards, 1% of N-Lauroylsarcosine (#L7414, Sigma-Aldrich) was added to the combined supernatant and rocked the mixture for 1 hour room temperature. After 1 hour, the mixture was spun for another hour at 100,000xg. The supernatant was saved as a soluble extract. The dark red brown pellet was then resuspended in 50mM Tris-HCl, pH = 7.5 at 0.2mL of mixture per gram of pellet, then stored at 4°C for further biochemical experiments.

### Biosensor analysis

#### Cell culture

“Biosensor” HEK293 cells stably expressing TauRD(P301S)-YFP were generated as previously described (32,34). Biosensor cells were maintained in a humidified 37°C, 5% CO2 incubator in Dulbecco’s modified Eagle medium (Gibco), and supplemented with 10% FBS with 1% penicillin/streptomycin (Gibco).

#### Brain lysate transfection

Tau brain lysate transfection was performed as previously described (32,34). In brief, Biosensor cells were plated at a density of 0.5X10^3^ cells per well in a 12-well plate (Catalog No.07-200-82, Corning). Twenty-four hours later, at ∼20∼25% confluency, cells were transduced with PS19 or control brain lysate. Transduction complexes were composed of 50 µL Opti-MEM (#31985070, ThermoFisher), 3 µL LT-1 transfection reagent (#MIR2020, Mirus Bio LLC), and 1 µL brain lysate (1ug/uL stock determined by BCA (5000001, Bio-Rad Protein Assay Kit I). After 24 hours, cells were fixed with 4% PFA/3% Sucrose fixation solution in PBS and imaged using ZEISS 710 confocal microscope.

#### Pharmacochemical modulation of tau aggregation

After 24 hours, culture media was replaced with fresh media with either PERK pathway inhibitors (GSK2656157, GSK2606414 or ISRIB) or modulators (Salubrinal or Selphin-1) for another 24 hours or more. Live cell imaging was performed using ZEISS 710 confocal microscope at 37°C, 5% CO_2_.

#### Tau aggregation in MEFs

*PERK^+/+^* or *PERK^-/-^*, and *eIF2α^S/S^* or *eIF2α^A/A^* MEF cells were transiently transduced with TauRD(P301S)-YFP plasmid using *TransIT* LT-1 Transfection reagent (#MIR2020, Mirus Bio LLC) and virally transduced by lab-prepared lentiviral infection following manufacturer’s instruction and incubated for 2 days. The cells were lysed with RIPA buffer, centrifuged at 14,000g for 10 min at 4°C; the supernatants saved for the western blot analysis. Cell imaging analysis was performed using ZEISS 710 confocal microscope.

### Preparation of Lentivirus

Lentivirus expressing TauRD(P301S)-YFP was prepared following Addgene lentivirus culture protocol. In brief, 293T packaging cells at 3×10^6^ cells per plate in DMEM complete media was plated in 10 cm cell culture dish. The TauRD(P301S)-YFP plasmid DNA with virus packing pDNAs (psPAX2; pMD2.G, Addgene) was transfected into 293T cells by 1mg/mL PEI transfection reagent and incubated for 36 hours. The resultant media was centrifuged at 150,000g for 90 min at 4°C. The concentrated virus was collected and stored at −80°C until further infection.

### Immunoblotting Analysis

*PERK^+/+^* and *PERK^-/-^*; *eIF2α^S/S^* and *eIF2α^A/A^* MEFs transduced with TauRD-YFP and Biosensor cell transfected with wild-type and PS19 brain lysates were lysed with SDS lysis buffer (2% SDS in PBS containing protease and phosphatase inhibitors (11836153001, Roche)) or RIPA buffer. Protein concentrations of the cell lysates were determined by BCA protein assay (Pierce). Equal amounts of protein were loaded onto 4–15% Mini-PROTEAN TGX precasted gels (Bio-Rad) and immunoblotted. The following antibodies and dilutions were used: anti-HT-7 at 1:1000, AT8 at 1:1000, YFP at 1:1000, HSP90 at 1:2000, Lamin A/C at 1:3000, GAPDH at 1:3000, T-PERK at 1:1000, P-PERK at 1:1000, eIF2*α* at 1:1000 and P-eIF2*α* at 1:1000. After overnight incubation with primary antibody, membranes were washed in TBS with 0.1% Tween-20, followed by incubation of a horseradish peroxidase-coupled secondary antibody (Cell Signaling). Immunoreactivity was detected using the SuperSignal West chemiluminescent substrate (Pierce) and BIO-RAD Universal Gel Molecular Imager.

### RNA-sequencing analysis

RNA-seq analysis was performed as previously described(36). In brief, frozen cells were collected, and RNA extraction, RNA quality control, and RNA sequencing were performed by BGI DNBSEQ Eukaryotic Strand-specific Transcriptome Resequencing (BGI); DNBSEQ stranded mRNA library, providing paired-end 100 bp reads at 30 million reads per sample. The sequencing data was filtered with SOAPnuke (v1.5.2)(66) and clean reads were mapped using HISAT2 (v2.0.4) to the reference genome (Homo_sapiens_GCF_000001405.38_GRCh38.p12 reference assembly)(67,68). The expression levels of the genes were calculated by RSEM (v1.2.12)(69). Differential expression analysis and statistical significance calculations between condition and experiment groups were assessed using R based software, DESeq2 (v 1.4.5)(70). The FDR q-value for the DESeq analysis is 0.1 (70). ER stress gene sets were collected from the results of the DESeq2 analysis and visualized using Prism GraphPad software. Violin plots comparing PS19 brain lysate transfection group and wild-type brain lysate transfection group were generated with the Log2 fold change data of the differential expression analysis.

### Human Alzheimer’s Disease brain RNA-Seq analysis

Human AD brain RNA sequencing datasets were collected from the NCBI gene expression omnibus (GEO) database: GSE173955(56) and GSE159699(55). Individual ER stress gene set of AD brains were extracted and normalized by non-AD control brain in each dataset and visualized using Prism GraphPad software. Violin plots comparing non-AD brain group and AD brain group were generated with the Log2 fold change data of the differential expression analysis.

### Gene Set Enrichment Analysis (GSEA) and Gene Ontology (GO) Analysis

gProfiler, a web-based application (https://biit.cs.ut.ee/gprofiler/), was used for GO pathway enrichment analysis. The gene sets from the bulk RNA sequencing analysis (Biosensor cell and human AD brain analysis: GSE173955(56) and GSE159699(55)) were entered, and Gene Ontology (GO) terms based on significant association were collected (p<0.05). GSEA (Gene Set Enrichment Analysis) software was downloaded (https://www.broadinstitute.org/gsea/) and also used to analyze the gene sets. Pre-ranked lists were entered with the same gene sets and ranked based on expression values relative to wild-type controls. Weighted analysis with the GO reference database was performed and GSEA enrichment plots presented.

### Statistical analysis

For RNA Seq gene expression data, differential gene expression analysis (DESeq) and the statistical significance (p-value) of differences between control and experimental groups (n=5 independent replicates) were assessed using R based software, DESeq2 (v 1.4.5)(70). The FDR q-value for the DESeq analysis is 0.1 (70); For the IRE1-, PERK-, ATF6-, and ERAD-regulated gene groups, the one sample t-test and Wilcoxon Signed Rank Test were used to calculate statistical significance of differences in the gene groups between control and experimental conditions. For individual genes, two-tailed Student’s t-test was used to calculate statistical significance of differences in gene expression between control and experimental conditions. For protein levels, two-tailed Student’s t-test was used to calculate the statistical significance of differences in protein levels between control and experimental conditions from immunoblot images captured by ImageJ and normalized to loading control images. For Biosensor cell aggregate puncta analysis, we used ImageJ to count cells and puncta visualized by confocal fluorescence microscopy and performed Student’s t-test and ANOVA Test followed by Tukey’s multiple comparisons test. The results were used with averages of all experiments ± standard deviation (SD). A probability of less than 0.05 was considered statistically significant and was annotated as *p≤0.05, **p≤0.01, ***p≤0.001, and ****p≤0.0001. All statistics were calculated using GraphPad Prism 9 software (GraphPad, San Diego, CA).

### Data availability

The RNA-Seq raw data and differential gene expression analyses from Biosensor cells are available under GEO accession number GSE217525.

## Acknowledgements

We thank Randal Kaufman for providing eIF2*α* and PERK MEF cell lines; Marc Diamond for providing Biosensor cells; Yusaku Nakabeppu for providing human AD brain tissue information; and Eun Jin Grace Lee and Luke Wiseman for helpful discussions and technical assistance.

## Author Contributions

G.P. and J.L. conceptualized and designed the project; G.P., L.C., K.X., and K.S. collected and analyzed bioinformatics database; G.P., L.C., K.X., L.S., N.H., S.P., and K.X. performed biochemistry analysis. G.P., S.P., H.M., and K.X. performed confocal imaging and data analysis; G.P., K.X., and H.M. designed and conducted all experiments using Biosensor cell image analysis. G.P., and J.L. co-wrote the manuscript. All authors read and approved the final manuscript.

## Funding and additional information

This work was supported by NIH R01NS088485 (J.L.), VA Merit I01RX002340 (J.L.), American Federation for Aging Research (J.L.), and Soonchunhyang University (Keon-Hyoung Song). The content is solely the responsibility of the authors and does not necessarily represent the official views of the National Institutes of Health.

## Abbreviations and nomenclature

EIF2AK3: Eukaryotic translation initiation factor 2 alpha kinase 3
PERK: Protein kinase R-like endoplasmic reticulum kinase
UPR: Unfolded Protein Response
ISR: Integrated Stress Response
ER: Endoplasmic Reticulum
AD: Alzheimer’s disease
PSP: Progressive Supranuclear Palsy
MAPT: Microtubule associated protein tau
WRS: Wolcott-Rallison syndrome
MEF: Mouse embryo fibroblast
YFP: Yellow fluorescence protein
GADD34: Growth arrest and DNA damage-inducible protein 34
PolyPhen-2: Polymorphism Phenotyping v2
PROVEAN: Protein Variation Effect Analyzer
SIFT: sorting intolerant from tolerant
CADD: Combined Annotation-Dependent Depletion

---

**Supplementary figure 1. Thapsigargin induces PERK-, IRE1-, ATF6 and ERAD-regulated gene expression in Biosensor cells.**

Biosensor cells were treated with thapsigargin (300nM) for 4 hours. The levels of PERK-, IRE1-, ATF6-, and ERAD-regulated genes were examined by RNA-Seq and normalized to gene expression levels in control (wild-type brain treated) cells. (**a, b**) Gene expression levels of 31 PERK-regulated genes in thapsigargin-treated cells relative to control are shown as log_2_ fold change. The graph (**a**) shows levels of the 5 PERK-regulated genes from figure 3e and *PERK* gene itself. The violin plot (**b**) shows levels of the entire PERK-regulated gene set. (**c, d**) Gene expression changes of 32 IRE1-regulated genes in thapsigargin-treated cells relative to control are shown. The graph (**c**) shows levels of the 5 IRE1-regulated genes most differentially expressed from figure 3g and *IRE1* gene itself. The violin plot (**d**) shows levels of the entire IRE1-regulated gene set. (**e, f**) Gene expression changes of 74 ATF6-regulated genes in thapsigargin-treated cells relative to control are shown. The graph (**e**) shows levels of the 5 ATF6-regulated genes most differentially expressed from figure 3i and *ATF6* gene itself. The violin plot (**f**) shows levels of the entire ATF6-regulated gene set. (**g**) Violin plot shows expression levels of 74 ERAD-regulated genes (drawn from ERAD pathway GO:0036593) in thapsigargin-treated cells relative to control. (**h**) GO analysis of RNA-Seq data from Biosensor cells treated with thapsigargin shows significant enrichment of ER stress and ERAD pathways in up-regulated genes, but not in down-regulated genes. (**i**) Wild-type brain, PS19 brain, or thapsigargin treated Biosensor cell protein lysates are immunoblotted for p-eIF2*α* or HSP90 (loading control). Error bars in **a**, **c**, and **e** represent mean ± standard deviation. Black squares represent 5 experimental replicates. (** *p*≤0.01, \*\*\*\**p*≤0.0001, not significant (ns), two-tailed Student’s *t-*test). The red horizontal line in **b**, **d**, **f, g** marks the median level of gene expression, and the thin horizontal blue lines delimit the upper and lower gene expression quartiles in the violin plots. (\*\*\*\**p*≤0.0001, one sample t-test and two-tailed Wilcoxon Signed Rank Test, n=5 experimental replicates). Detailed gene expression RNA-Seq information is shown in Supplemental data 3 and 4.

**Supplementary figure 2. Gene Set Enrichment Analysis (GSEA) of ER stress and ERAD in Biosensor cells after PS19 brain or thapsigargin treatment. a,** GSEA plot of ER stress genes in PS19 treated Biosensor cells relative to wild-type brain treated cells. **b**, GSEA plot of ERAD genes in PS19 treated Biosensor cells relative to wild-type brain treated cells. **c**, GSEA plot of ER stress genes in thapsigargin treated Biosensor cells relative to wild-type brain treated cells. **d**, GSEA plot of thapsigargin treated Biosensor cells relative to wild-type brain treated cells. The Normalized Enrichment Score (NES), Nominal p-value, and False Discovery Rate (FDR) q-value for each GSEA plot are shown.

