## Supplementary figures and images for "Neurodegeneration Risk Factor, *EIF2AK3* (*PERK*), Influences Tau Protein Aggregation"

### Supplemental figures

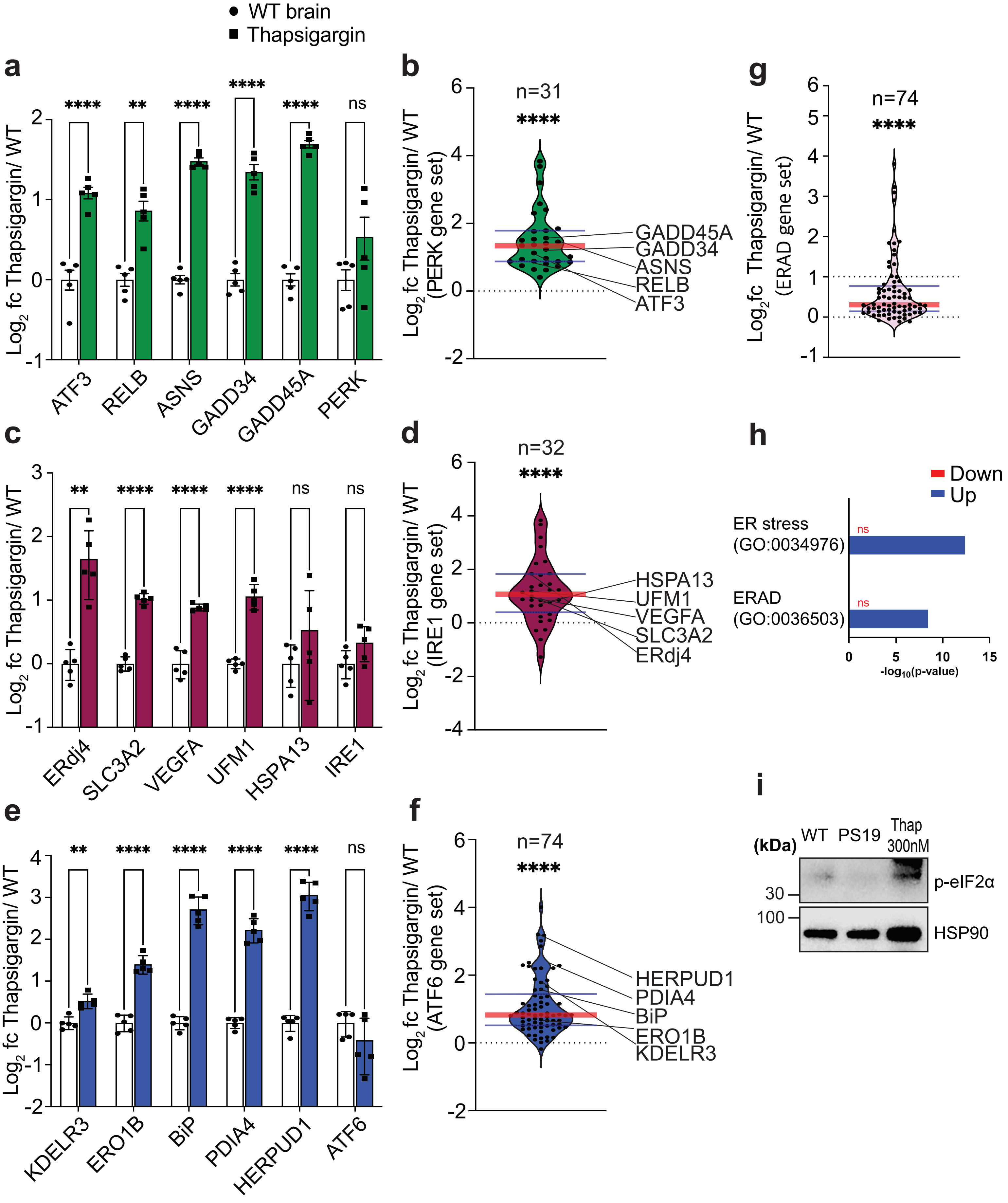


Suppl. Figure 1

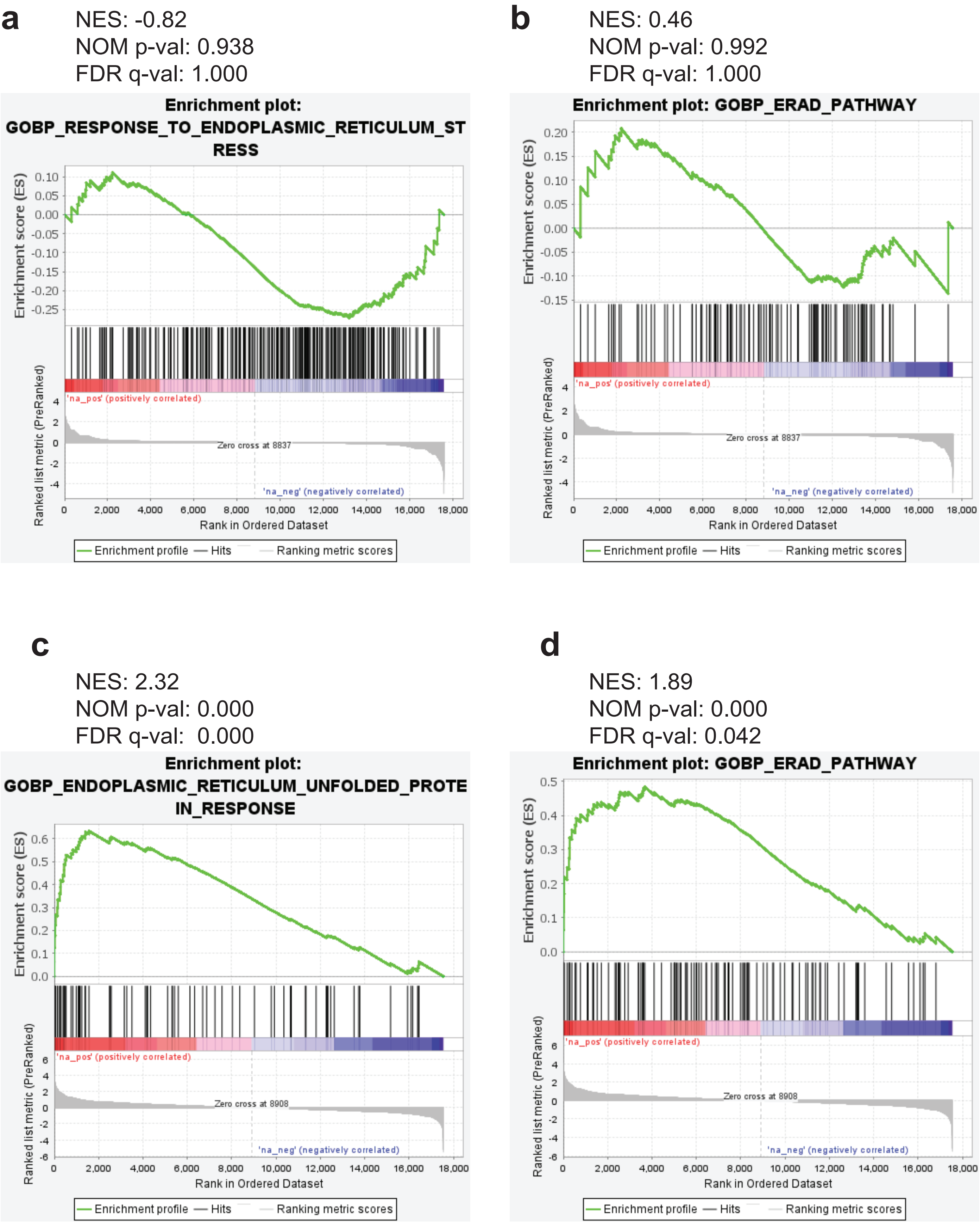


Suppl. Figure 2
